# Discovery and Validation of *RUNX1* DNA Methylation in Differentiating Papillary Thyroid Cancer from Benign Nodules

**DOI:** 10.1101/2023.04.10.536270

**Authors:** Junjie Li, Yifei Yin, Haixia Huang, Mengxia Li, Hong Li, Minmin Zhang, Chenxia Jiang, Rongxi Yang

**Affiliations:** Department of Epidemiology and Biostatistics, School of Public Health, Nanjing Medical University, 210000 Nanjing, China; Department of Thyroid and Breast Surgery, The Affiliated Huai’an Hospital of Xuzhou Medical University and The Second People’s Hospital of Huai’an; Department of Pathology, The Affiliated Huai’an Hospital of Xuzhou Medical University and The Second People’s Hospital of Huai’an; Department of Pathology, The Affiliated Hospital of Nantong University, 226001 Nantong, China

**Keywords:** Benign thyroid nodule, Papillary thyroid carcinoma, DNA methylation, *RUNX1*, Biomarker

## Abstract

Although most thyroid nodules can be diagnosed preoperatively by thyroid ultrasonography and fine-needle aspiration biopsy, it remains a challenge to accurately identify malignancy of thyroid nodules when the biopsy is indeterminate. This study aims to explore a novel biomarker to distinguish benign and malignant thyroid nodules. Tissue samples from patients with Stage I&II papillary thyroid carcinoma (PTC) and benign thyroid nodules (BTN) were collected for genome profiling by methylation EPIC 850K array and RNA-Sequencing. Genes with significantly differential DNA methylation and inverse mRNA expression were filtered out. The altered methylation of *RUNX1* gene was validated in two independent case-control studies with a total of 699 formalin fixed paraffin-embedded (FFPE) samples using mass spectrometry and calculated by binary logistic regression analysis. Hypomethylation of *RUNX1* gene in PTC patients compared to BTN subjects was verified in Validation Ⅰ (140 PTC vs. 189 BTN, ORs ≥ 1.50 per-10% methylation, *P* ≤ 4.40E-05, for all measurable CpG sites) and Validation Ⅱ (184 PTC vs. 186 BTN, ORs ≥ 1.72 per-10% methylation, *P* ≤ 2.38E-11, for all measurable CpG sites). Besides, *RUNX1* methylation achieved good accuracy in differentiating early-stage PTC from BTN in Validation Ⅰ (AUC: 0.74) and Validation Ⅱ (AUC: 0.79). Gender- and age-stratified analysis revealed *RUNX1* hypomethylation as an important risk factor for thyroid disease in younger women. We disclosed a significant association between *RUNX1* hypomethylation and PTC, suggesting *RUNX1* methylation based on FFPE tissue samples as a potential biomarker for predicting malignancy of thyroid nodules.

## Introduction

Thyroid cancer (TC) is the most common endocrine malignancy[1, 2]. Papillary thyroid cancer (PTC) with low malignancy and good prognosis is the most common subtype, accounting for about 85% of all TC cases[3]. TC is most common in younger women, and its incidence varies by region, race, age and gender[1, 4]. In recent years, the incidence of TC has increased annually by 20% in some Asian countries, including South Korea, Japan and China[5]. According to the global statistics in 2020, the incidence of TC ranks ninth among the malignant cancers worldwide. In China, the incidence of TC ranks the fourth among women and the second among younger people, posing a serious burden on health care and social economy[6].

The thyroid gland is small and has abundant peripheral blood vessels. Diagnosis of thyroid nodules aims to accurately identify malignancy and avoid overtreatment. Fine needle aspiration biopsy (FNAB) is used as the principal method for preoperative diagnosis of TC in clinical practice, which has the advantages of little damage to the body and avoiding tumor cell dissemination[7]. The Bethesda System for Reporting Thyroid Cytopathology categorizes the FNAB results into six groups (Ⅰ-Ⅵ) according to the cytological features[8]. Among them, the Bethesda Ⅲ-Ⅴ nodules were indeterminate nodules, which account for about 20% and need further evaluation[9]. As the accuracy of FNAB results largely depends on the quality of embedding and biopsy staining as well as the experiences of the pathologists, the sensitivity and specificity of FNAB was variant from 54% to 90% and from 60% to 98% by different clinical centers [10, 11]. High-quality histopathology is essential for diagnosis and treatment of patients. However, current health statistics in some developing countries point to a serious shortage of clinical pathologists, and the professional proficiency of doctors is not uniform[12, 13]. Molecule pathological techniques could be more objective and thus have also been developed, including Afirma^®^ Genomic Sequencing Classifier, ThyroSeq^®^ v3, ThyGenX/ThyraMIR, and RosettaGX Reveal, which detect a variety of changes at gene rearrangements, genomic mutations, mRNA and microRNA expression[14, 15]. Nevertheless, these tests are always challenged by the instability of RNA samples and shared mutations between malignant and benign thyroid tissues, and their clinical application is limited. Hence, it has great importance to explore an objective, reliable and stable biomarker to differentiate benign and malignant thyroid nodules.

DNA methylation is a common type of epigenetic modification, which can influence gene expression by regulating gene transcription[16]. Numerous studies have shown that abnormal DNA methylation can lead to diseases (e.g., tumors)[17]. During tumorigenesis, oncogenes are often activated by promoter hypomethylation[18]. DNA methylation alterations occur in the early stage of tumor formation and are involved during the whole malignant progression of various cancers. DNA methylation has the advantages of high tissue specificity, high stability, as well as mature and reliable quantitative detection technology[19]. In addition, the collection, processing and storage of clinical samples are easy to operate, and the amount of sample required for detection is small. Therefore, DNA methylation has been actively investigated and widely applied in clinical practice as an emerging biomarker for cancer diagnosis.

Previous reports on DNA methylation-based differential diagnosis of thyroid nodules mostly focused on DNA methylation of known oncogenes (e.g., *RASSF1* and *SLC5A8*)[20, 21]. However, due to low sensitivity and specificity of diagnosis, insufficient sample size and lack of external validation studies, these studies have not achieved satisfactory results. Here, to discover and verify the abnormal DNA methylation signatures in early-stage PTC cases compared to BTN patients, we performed 850K beadchip array and RNA-sequencing in tissue samples from 17 BTN subjects and 15 early-stage PTC cases. The selected PTC-related gene with significant methylation changes was further validated by mass spectrometry in two independent case-control studies, aiming to identify potential biomarkers for differentiating benign and malignant thyroid tumors.

## Materials and methods

### Study design and patients

In the discovery study and two independent validation studies, the inclusion criteria for malignant nodules include: (1) early-stage (stage Ⅰ & Ⅱ) PTC with classic papillary structures; (2) no distant metastasis or other simultaneous cancers; (3) no related therapy. BTN patients were matched to PTC cases by age, gender and the year of diagnosis. All cases were histopathologically diagnosed by two qualified pathologists, and clinical TNM stage of each malignant case was determined on the basis of the 8^th^ edition American Joint Committee on Cancer staging system[22].

In the discovery study, a total of 32 fresh frozen tissue samples were collected from the Affiliated Huai’an Hospital of Xuzhou Medical University from 2019 to 2020, with 15 early-stage PTC cases and 17 BTN subjects, respectively. All the subjects were women and the age at diagnosis ranged from 23 to 76 years old.

Two independent validation studies with formalin-fixed and paraffin-embedded (FFPE) tissue samples were conducted for validation. Validation Ⅰ: 140 early-stage PTC cases and 189 age- and gender-matched BTN patients were collected from the Affiliated Huai’an Hospital of Xuzhou Medical University from 2019 to 2020. Women accounted for 77.14% (108/140) in PTC cases and 78.84% (149/189) in BTN groups. The median ages and interquartile ranges (IQRs) in PTC and BTN groups were 50.00 (43.00-57.00) and 53.00 (46.00-60.50) years old, respectively. Validation Ⅱ: a total of 184 early-stage PTC cases and 186 age- and gender-matched BTN subjects were collected from the Affiliated Hospital of Nantong University in 2020. Women accounted for 78.80% (145/184) in PTC and 77.42% (144/186) in BTN groups. The median ages and interquartile ranges (IQRs) in PTC and BTN groups were 49.00 (33.25-56.00) and 49.50 (37.00-55.25) years old, respectively. The detailed clinical characteristics of the participants, including tumor length, tumor size, involved lymph node and tumor stage were shown in Table 1.

**Table 1.**
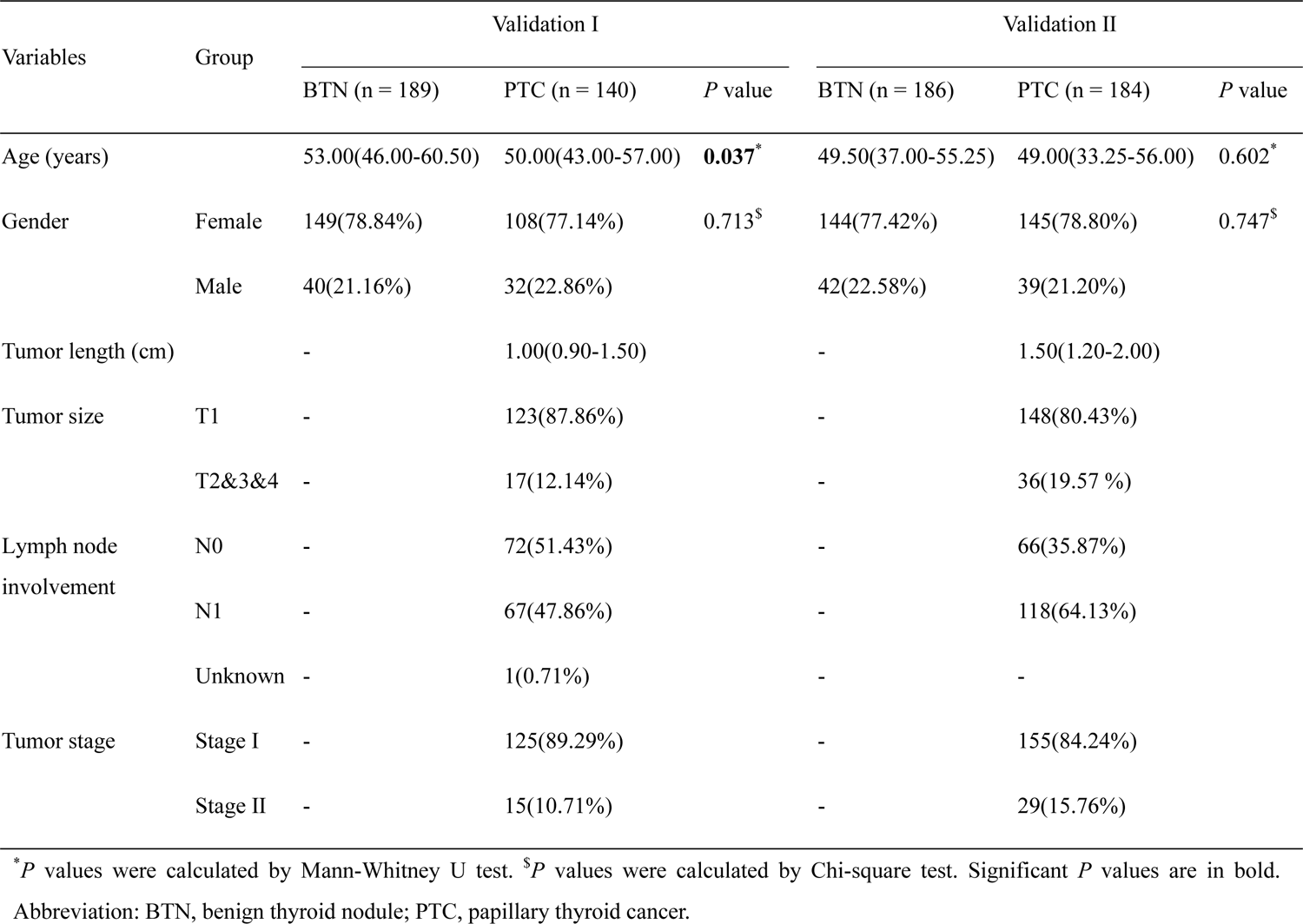
Clinical characteristics of the participants

The studies were approved by the Ethics Committee of the Nanjing Medical University, the Affiliated Huai’an Hospital of Xuzhou Medical University and the Affiliated Hospital of Nantong University. Informed consent was obtained from all patients.

### Illumina Infinium Human Methylation EPIC 850K beadchip array and RNA-Sequencing

The study design and flow chart were shown in Figure 1. In the discovery study, according to the manufacturer’s instructions, genomic DNA and total RNA were isolated from 32 fresh frozen tissue samples using a FastPure Blood/Cell/Tissue/Bacteria DNA Isolation Mini Kit (DC112, Vazyme, Nanjing, China) and a FastPure Cell/Tissue Total RNA Isolation Kit (RC101, Vazyme, Nanjing, China), respectively. Genome-wide DNA methylation profiles were analyzed by Illumina Infinium Human Methylation EPIC 850K beadchip array with single-nucleotide resolution. Probes meeting the following criteria were considered as differentially methylated: (1) methylation difference (|delta beta|) between BTN and PTC groups ≥ 0.2; (2) an FDR-corrected *P* value < 0.01; (3) no adjacent single nucleotide polymorphisms (SNPs); and (4) on the promoter region or the 1st exon of gene body. At the same time, the next generation RNA sequencing technology was used to detect the expression levels of mRNA. Genes mRNA expression with |fold change| ≥ 2 and an FDR-corrected *P* value < 0.05 were thought to be differentially expressed. Through the intersection of 850K beadchip array results and RNA sequencing results, genes with significant differences in methylation and mRNA expression were screened out.

**Figure 1.**
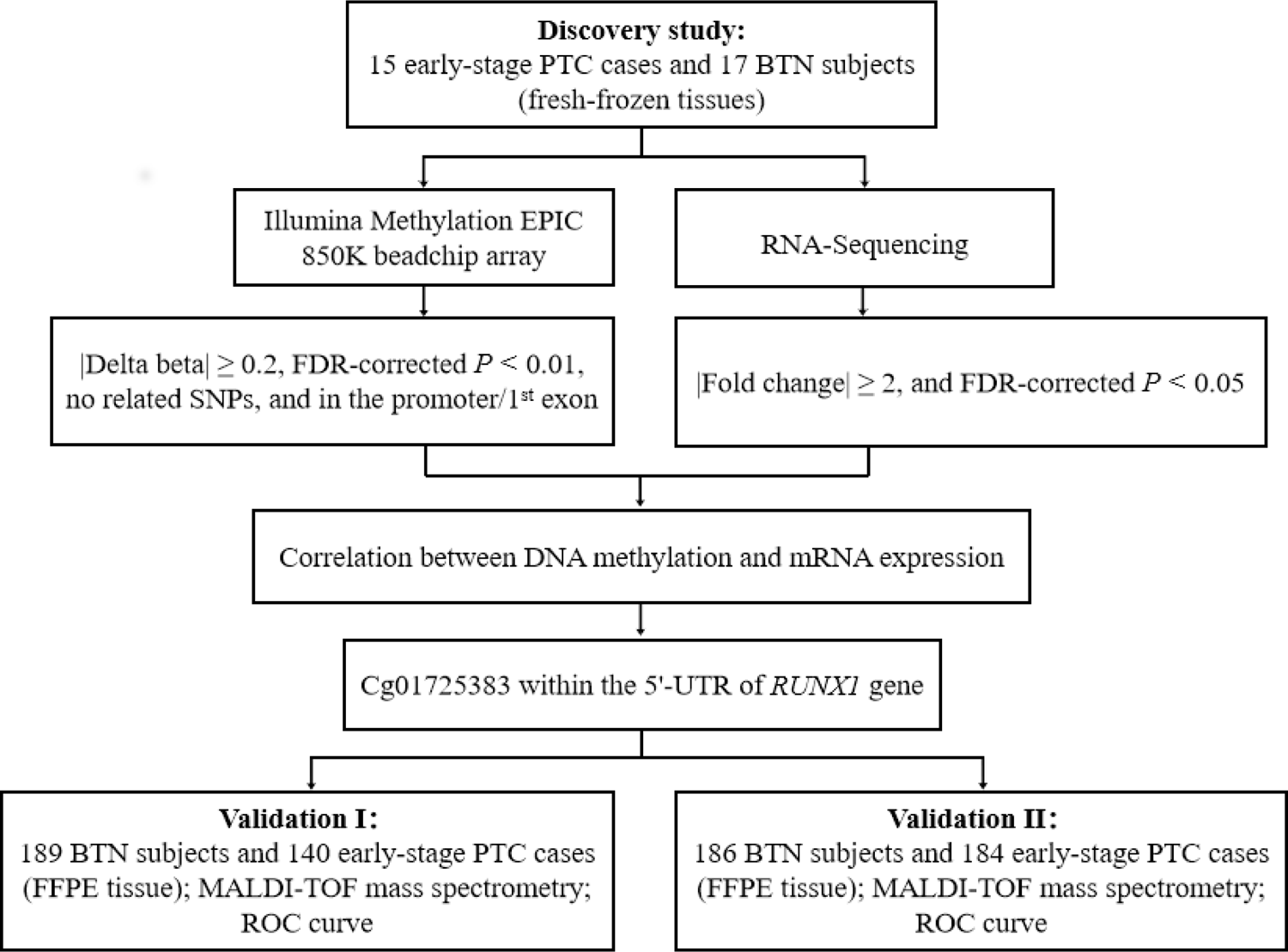
Study design and flow chart. The 32 fresh-frozen tissue samples in the discovery study were subjected to Illumina Methylation EPIC 850K beadchip array and RNA-Sequencing. The cg01725383 within 5’-UTR of *RUNX1* gene were selected by comprehensive analysis of binary-omics data in a stepwise selection manner. Further validations with FFPE tissue samples were carried out in our two independent case-control studies (Validation Ⅰ and Validation Ⅱ). Abbreviation: PTC, papillary thyroid cancer; BTN, benign thyroid nodule; FFPE, formalin-fixed and paraffin-embedded; SNP, single nucleotide polymorphism; UTR, untranslated region; MALDI-TOF, matrix-assisted laser desorption ionization time-of-flight; ROC, receiver operating characteristic.

### Matrix-assisted laser desorption Ionization time-of-flight (MALDI-TOF) mass spectrometry

By the UCSC Genome Browser, a 196 bp amplicon (Chr21: 36,259,768 - 36,259,964 build GRCh37/hg19) at the first Exon of *RUNX1* gene and covering 4 CpG sites was designed (Figure 2). The whole gene sequence of *RUNX1* amplicon was GGAACTTCCAAATAATGTGTTTGCTGAT**CG**TTTTACTCTT**CG**CATAAATATTTTAGGAAGTGT ATGAGAATTTTGCCTTCAGGAACTTTTCTAACAGCCAAAGACAGAACTTAACCTCTGCAAG CAAGATT**CG**TGGAAGATAGTCTCCACTTTTTAATGCACTAAGCAAT**<u>CG</u>**GTTGCTAGGAGCC CATCCTGGGTC, and the target sequence was amplified by polymerase chain reaction (PCR). Forward primer: aggaagagagGGAATTTTTAAATAATGTGTTTGTTGAT; Reverse primer: cagtaatacgactcactatagggagaaggctAACCCAAAATAAACTCCTAACAACC. Upper case letters indicated the sequence specific primer regions, and non-specific tags were shown in lower case letters. The EpiTyper assay detected the methylation levels of four CpG sites and yielded four distinguishable mass peaks. There were no single nucleotide polymorphisms (SNPs) located in the primer regions or overlapped with any of the CpG sites in *RUNX1* amplicon. The measurable CpG sites are in bold. Cg01725383 is underlined.

**Figure 2.**
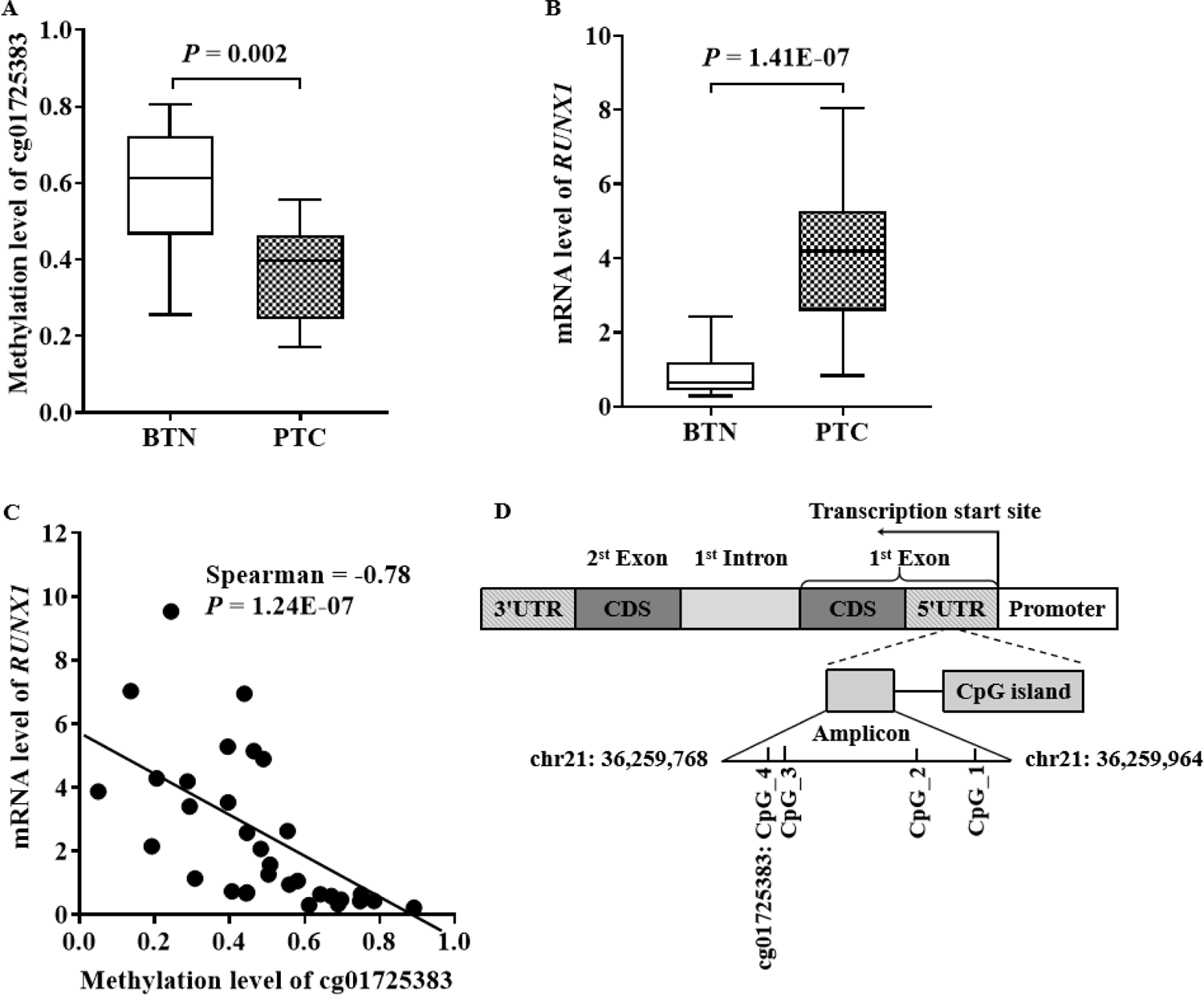
Discovery of differentially methylated cg01725383 in *RUNX1* gene in fresh-frozen tissue samples in the discovery study. (A-B) Box plots for DNA methylation levels of cg01725383 detected by 850K beadchip array (A) and for mRNA expression levels of *RUNX1* measured by RNA-Sequencing (B) in BTN subjects and PTC cases. (C) The correlation between methylation levels of cg01725383 and *RUNX1* mRNA expression. (D) Schematic diagram of the target amplicon within *RUNX1* gene. A 196 bp amplicon covering four differentially methylated CpG sites was located at 5’-UTR of the *RUNX1* gene (Chr21: 35,259,768 – 36,259,964, build GRCh37/hg19, defined by the UCSC Genome Browser).

MALDI-TOF mass spectrometry was used for semi-quantitative analysis of DNA methylation levels. All the PTC and BTN samples were processed in parallel. First, DNA was extracted from the FFPE tissue samples using FastPure FFPE DNA Isolation Kit (DC105, Vazyme, Nanjing, China). Next, DNA was treated with sodium bisulfite using EZ-96 DNA Methylation Gold Kit (Zymo Research, Orange, USA), through which non-methylated cytosine (C) in CpG site was converted to uracil (U), whereas methylated cytosine remained unchanged. Finally, CpG methylation levels of FFPE tissue samples were measured by MALDI-TOF mass spectrometry, as described in our previous study[23]. In brief, the target sequence was amplified by polymerase chain reaction. Then the amplification products were treated with Shrimp Alkaline Phosphatase and followed by T-cleavage using RNase A. After cleaning residual ions with resin, methylation level of each sample was quantified and collected by the MassARRAY system.

### Statistical analyses

All statistical analyses were performed by IBM SPSS Statistics 25.0 version and GraphPad Prism (version 9.0). Nonparametric tests including Mann-Whitney U test and Chi-square test were applied to compare the difference between PTC and BTN groups, and Kruskal-Wallis test was used to compare the methylation levels of clinical characteristics between multiple groups. Binary logistic regression analysis was conducted to calculate odds ratios (ORs) per −10% methylation and their 95% confidence intervals (95% CIs) after adjusting for age, gender. Spearman correlation was used to determine the relationship between variables. Receiver operating characteristic (ROC) curve was used to evaluate goodness of fit. All tests were two-sided, and a *P* value less than 0.05 was considered to be statistically significant.

### Data and Reagent Availability Statement

The authors affirm that all data and reagent necessary for confirming the conclusions of the article are represented fully within the article, figures, and tables.

## Results

### Discovery of *RUNX1* hypomethylation to distinguish early-stage PTC cases from BTN patients

In the discovery study, we performed epigenome-wide screening using 850K beadchip array and whole transcriptome screening using RNA-Sequencing in parallel in the fresh frozen tissue samples of 15 early-stage PTC and 17 BTN. By overlapping differentially methylated and expressed genes, a CpG site, cg01725383, located at the 5’ untranslated region (5’ - UTR) of *RUNX1* gene was identified (Figure 1). Compared to BTN subjects, the CpG site cg01725383 showed significant hypomethylation in early-stage PTC cases (median methylation value: 0.61 in BTN and 0.40 in PTC, FDR-corrected *P* value = 0.002) (Figure 2A). It is well known that the alteration in DNA methylation could affect gene expression. Indeed, compared to BTN subjects, the mRNA levels of *RUNX1* in PTC cases were significantly increased (median mRNA expression value: 0.64 in BTN and 4.19 in PTC, FDR-corrected *P* value = 1.41E-07) (Figure 2B). Moreover, there was a significant inverse correlation between the methylation levels of cg01725383 and the mRNA levels of *RUNX1*, with the Spearman’s correlation coefficient of −0.78 (*P* value = 1.24E-07) (Figure 2C). A 196-bp amplicon containing the cg01725383 site and flanking CpG sites was designed for further validation by MALDI-TOF mass spectrometry (Figure 2D). No known SNPs or CpG sites were found within the primer sequence. Within the amplicon, the EpiTyper system measured the methylation levels of four CpG sites and produced four distinguishable mass peaks. Cg01725383 was referred as CpG_1.

### Validation of the association between *RUNX1* hypomethylation and PTC in FFPE tissues by two independent case-control studies

The validation of the *RUNX1* methylation difference between early-stage PTC cases and BTN subjects was carried out with FFPE tissue samples in two independent case-control studies (Validation Ⅰ and Validation Ⅱ) (Figure 1).

In Validation I consisting of 140 early-stage PTC cases and 189 age- and gender-matched BTN subjects, CpG_1 presented the most significantly reduced methylation levels in PTC cases compared to BTN subjects (methylation values of BTN and PTC: 0.41 vs. 0.25). Similarly, CpG_2, CpG_3 and CpG_4 were hypomethylated in PTC cases than those in BTN subjects (methylation values of BTN and PTC, respectively: 0.39 vs. 0.27; 0.08 vs. 0.03; 0.15 vs. 0.08) (Figure 3A and Table 2). Furthermore, there was a significant association between *RUNX1* hypomethylation and early-stage PTC. After adjusting for age and gender, the ORs per 10% reduced methylation of all CpG sites in *RUNX1* amplicon ranged from 1.50 to 1.96 (all *P* values ≤ 4.40E-05; Table 2).

**Figure 3.**
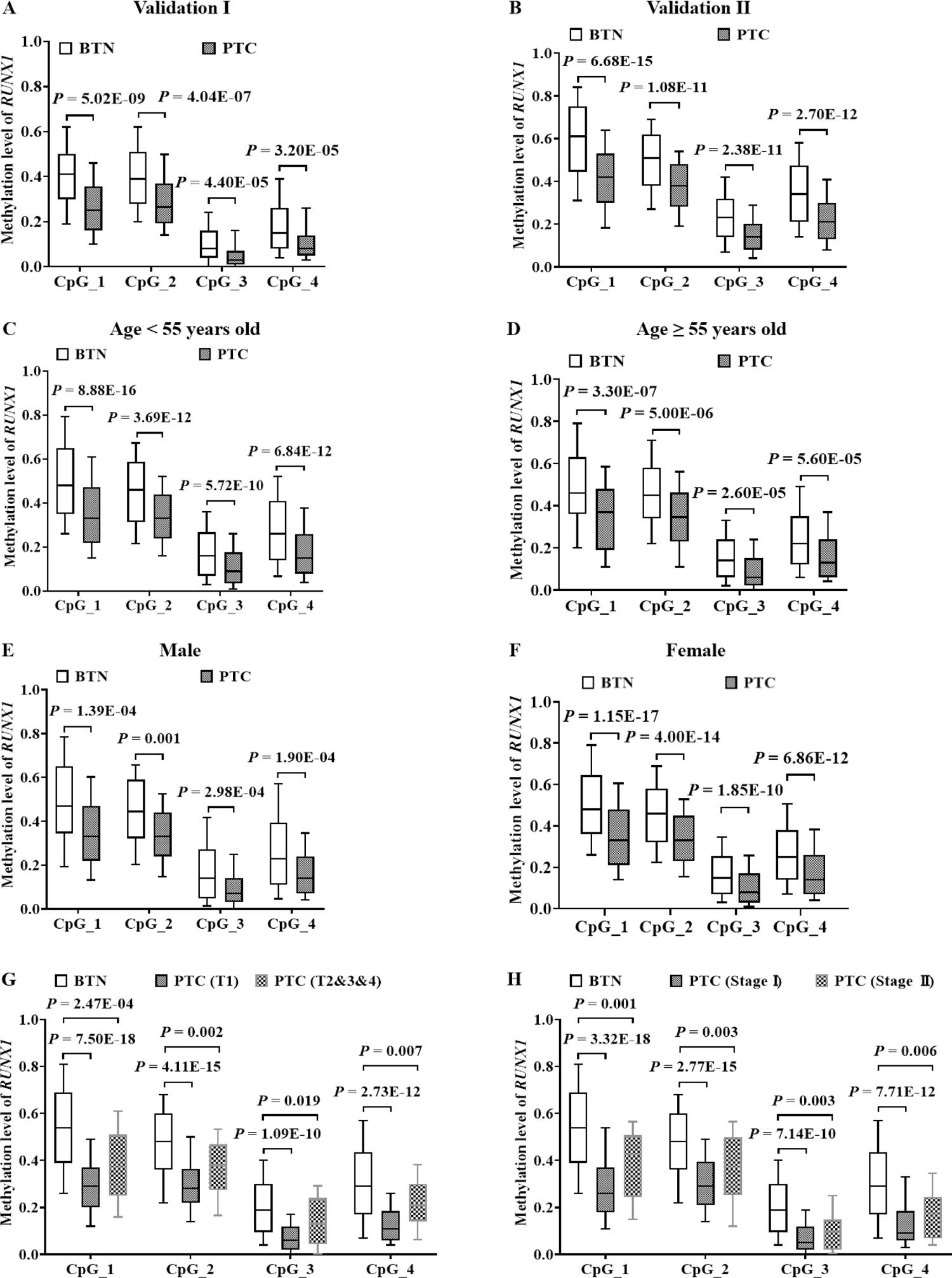
Validation of *RUNX1* hypomethylation in PTC cases compared to BTN subjects in two independent case-control studies, and discovery the influence of age, gender, tumor size and tumor stage on PTC-associated *RUNX1* hypomethylation. (A-B) Box plots for DNA methylation levels of the four CpG sites in *RUNX1* amplicon in BTN and PTC detected by mass spectrometry in Validation Ⅰ (A) and Validation Ⅱ (B). Furthermore, Validation Ⅰ and Validation Ⅱ were combined and stratified by the age of 55 years old, gender, tumor size or tumor stage, respectively. (C-H) Box plots for DNA methylation levels of the four CpG sites in *RUNX1* amplicon in BTN and PTC detected by mass spectrometry in age < 55 years old group (C), age ≥ 55 years old group (D), male group (E), female group (F), tumor size (G) and tumor stage (H), respectively. All *P* values were calculated by logistic regression with covariates-adjusted.

**Table 2.** The association between *RUNX1* methylation and early-stage PTC in two validation studies

| CpG sites | BTN<br>median (IQR) | PTC<br>median (IQR) | OR (95% CI) per | *P* value<sup>s</sup> |
| --- | --- | --- | --- | --- |
| **Validation I (189 BTN vs. 140 PTC)** |  |  |  |  |
| CpG_1 | 0.41(0.30-0.50) | 0.25(0.16-0.36) | 1.63(1.39-1.92) | **5.02E-09** |
| CpG_2 | 0.39(0.28-0.51) | 0.27(0.19-0.37) | 1.50(1.28-1.76) | **4.04E-07** |
| CpG_3 | 0.08(0.04-0.16) | 0.03(0.01-0.07) | 1.96(1.42-2.70) | **4.40E-05** |
| CpG_4 | 0.15(0.08-0.26) | 0.08(0.05-0.14) | 1.56(1.26-1.92) | **3.20E-05** |
| **Validation II (186 BTN vs. 184 PTC)** |  |  |  |  |
| CpG_1 | 0.62(0.45-0.75) | 0.41(0.30-0.53) | 1.72(1.50-1.97) | **6.68E-15** |
| CpG_2 | 0.52(0.38-0.62) | 0.38(0.28-0.48) | 1.72(1.47-2.01) | **1.08E-11** |
| CpG_3 | 0.23(0.14-0.32) | 0.14(0.08-0.20) | 2.13(1.71-2.67) | **2.38E-11** |
| CpG_4 | 0.35(0.22-0.48) | 0.21(0.12-0.30) | 1.79(1.52-2.11) | **2.70E-12** |

<sup>s</sup>Logistic regression with adjustment for age and gender. Significant *P* values are in bold.
Abbreviation: BTN, benign thyroid nodule; PTC, papillary thyroid cancer; IQR, interquartile range.

Consistent results were obtained in Validation II consisting of 184 early-stage PTC cases and 186 age- and gender-matched BTN subjects. We observed that the *RUNX1* gene was significant hypomethylated in PTC cases compared to BTN subjects. CpG_1 exhibited the most significantly reduced methylation levels in PTC cases compared to BTN subjects (methylation values of BTN and PTC: 0.62 vs. 0.41). Similarly, CpG_2, CpG_3 and CpG_4 were hypomethylated in PTC cases than those in BTN subjects (methylation values of BTN and PTC respectively: 0.52 vs. 0.38; 0.23 vs. 0.14; 0.35 vs. 0.21) (Figure 3B and Table 2). Binary logistic regression analyses revealed that hypomethylation of *RUNX1* had robust associations with early-stage PTC. Covariates-adjusted ORs ranged from 1.72 to 2.13 per 10% reduction in methylation of each *RUNX1* CpG site (all *P* values ≤ 2.38E-11; Table 2).

### Combination analysis of the association between FFPE tissue-based *RUNX1* hypomethylation and early-stage PTC stratified by age and gender

We next investigated whether the association between *RUNX1* hypomethylation and early-stage PTC was influenced by age and gender. To avoid the possible bias due to small sample size, the subjects in two validations were combined (324 PTC cases vs. 375 BTN subjects) and binary logistic regression analysis was performed in gender- or age-stratified groups.

All participants were stratified by the age of 55 years old, which is a cut-off threshold for tumor staging of PTC cases[22]. For age < 55 years old group, all the four CpG sites in *RUNX1* amplicon showed significantly decreased methylation levels in PTC cases compared to BTN subjects (Figure 3C and Table 3). After adjusting for gender, the ORs per 10% reduced methylation of the four CpG sites ranged from 1.62 to 1.89 (all *P* values ≤ 5.72E-10; Table 3). Consistently, for age ≥ 55 years old group, the four CpG sites were significantly hypomethylated in PTC cases than in BTN subjects (Figure 3D and Table 3). After adjusting for gender, the ORs per 10% reduced methylation of the four CpG sites ranged from 1.51 to 1.86 (all *P* values ≤ 5.60E-05; Table 3). Compared with age ≥ 55 years old group, *RUNX1* methylation and OR values showed a greater difference between BTN subjects and PTC cases in age < 55 years old group, indicating that the *RUNX1* hypomethylation was a great risk factor for younger people.

**Table 3.** The association between *RUNX1* methylation and early-stage PTC stratified by age or gender after combining Validation Ⅰ and Validation Ⅱ

| CpG sites | BTN<br>median (IQR) | PTC<br>median (IQR) | OR (95% CI) per | *P* value<sup>s</sup> |
| --- | --- | --- | --- | --- |
| **< 55 years old (236 BTN vs. 226 PTC)** |  |  |  |  |
| CpG_1 | 0.48(0.35-0.65) | 0.33(0.22-0.47) | 1.68(1.48-1.90) | **8.88E-16** |
| CpG_2 | 0.46(0.31-0.59) | 0.33(0.24-0.44) | 1.62(1.42-1.86) | **3.69E-12** |
| CpG_3 | 0.16(0.07-0.27) | 0.09(0.04-0.18) | 1.89(1.55-2.31) | **5.72E-10** |
| CpG_4 | 0.26(0.14-0.41) | 0.15(0.08-0.26) | 1.69(1.46-1.97) | **6.84E-12** |
| **≥ 55 years old (139 BTN vs. 98 PTC)** |  |  |  |  |
| CpG_1 | 0.46(0.36-0.63) | 0.37(0.19-0.48) | 1.53(1.30-1.81) | **3.30E-07** |
| CpG_2 | 0.45(0.34-0.58) | 0.35(0.23-0.46) | 1.51(1.26-1.80) | **5.00E-06** |
| CpG_3 | 0.14(0.06-0.24) | 0.06(0.02-0.15) | 1.86(1.39-2.49) | **2.60E-05** |
| CpG_4 | 0.22(0.12-0.35) | 0.13(0.06-0.24) | 1.52(1.24-1.87) | **5.60E-05** |
| **Male (82 BTN vs. 71 PTC)** |  |  |  |  |
| CpG_1 | 0.47(0.35-0.65) | 0.33(0.22-0.47) | 1.49(1.21-1.82) | **1.39E-04** |
| CpG_2 | 0.45(0.32-0.59) | 0.33(0.24-0.44) | 1.44(1.16-1.80) | **0.001** |
| CpG_3 | 0.14(0.05-0.27) | 0.07(0.03-0.14) | 1.86(1.33-2.60) | **2.98E-04** |
| CpG_4 | 0.23(0.11-0.40) | 0.14(0.07-0.24) | 1.63(1.26-2.11) | **1.90E-04** |
| **Female (293 BTN vs. 253 PTC)** |  |  |  |  |
| CpG_1 | 0.48(0.36-0.65) | 0.33(0.21-0.48) | 1.65(1.47-1.85) | **1.15E-17** |
| CpG_2 | 0.46(0.32-0.58) | 0.33(0.23-0.45) | 1.62(1.43-1.83) | **4.00E-14** |
| CpG_3 | 0.15(0.07-0.26) | 0.08(0.03-0.17) | 1.85(1.53-2.23) | **1.85E-10** |
| CpG_4 | 0.25(0.14-0.38) | 0.14(0.07-0.26) | 1.61(1.41-1.85) | **6.86E-12** |

<sup>s</sup>Logistic regression with adjustment for covariates. Significant *P* values are in bold.
Abbreviation: BTN, benign thyroid nodule; PTC, papillary thyroid cancer; IQR, interquartile range.

When stratified by gender, for males, all the four CpG sites in *RUNX1* amplicon in PTC cases showed significantly decreased methylation levels compared to BTN subjects (Figure 3E and Table 3). After adjusting for age, the ORs per 10% reduced methylation of the four CpG sites ranged from 1.44 to 1.86 (all *P* values ≤ 0.001; Table 3). Similarly, for females, the four CpG sites were significantly hypomethylated in PTC cases than in BTN subjects (Figure 3F and Table 3). After adjusting for age, the ORs per 10% reduced methylation of the four CpG sites ranged from 1.61 to 1.85 (all *P* values ≤ 1.85E-10; Table 3). Compared with male group, *RUNX1* methylation levels and OR values showed a greater difference between BTN subjects and PTC cases in female group, indicating that the *RUNX1* hypomethylation was a great risk factor for female.

### Combination analysis of the correlation between *RUNX1* methylation and the clinical characteristics of early-stage PTC

Next, 324 PTC cases combining two validations were stratified by clinical characteristics, and *RUNX1* methylation differences were analyzed between subgroups by nonparametric tests. There was no significant correlation between *RUNX1* methylation and tumor length, tumor size, lymph node involvement or tumor stage (all *P* values > 0.05; Table 5). In spite of that, all the four CpG sites in *RUNX1* amplicon showed significantly decreased methylation levels in PTC cases with T1 or at Stage Ⅰ compared to BTN subjects (Figure 3G, 3H and Table 4). After adjusting for age and gender, the ORs per 10% reduced methylation of the four CpG sites ranged from 1.50 to 1.66 for PTC cases with T1 and 1.48 to 1.59 for PTC cases at Stage Ⅰ (all *P* values ≤ 7.14E-10; Table 4). All above results suggested that *RUNX1* methylation is more effective in differentiating early-stage PTC cases with T1 or at Stage Ⅰ from BTN patients, although the accuracy of this conclusion might be limited by small sample size of PTC cases with T2&3&4 or at Stage II.

**Table 4.** The association between *RUNX1* methylation and early-stage PTC stratified by tumor size or tumor stage after combining Validation Ⅰ and Validation Ⅱ

| CpG sites | BTN<br>median (IQR) | PTC<br>median (IQR) | OR (95% CI) per | *P* value<sup>s</sup> |
| --- | --- | --- | --- | --- |
| **375 BTN vs. 271 PTC (T1)** |  |  |  |  |
| CpG_1 | 0.48(0.36-0.65) | 0.33(0.21-0.46) | 1.50(1.37-1.65) | **7.50E-18** |
| CpG_2 | 0.46(0.32-0.58) | 0.32(0.23-0.44) | 1.52(1.37-1.69) | **4.11E-15** |
| CpG_3 | 0.15(0.06-0.26) | 0.08(0.03-0.16) | 1.66(1.42-1.93) | **1.09E-10** |
| CpG_4 | 0.25(0.13-0.39) | 0.14(0.07-0.24) | 1.50(1.34-1.68) | **2.73E-12** |
| **375 BTN vs. 53 PTC (T2&3&4)** |  |  |  |  |
| CpG_1 | 0.48(0.36-0.65) | 0.37(0.25-0.51) | 1.34(1.15-1.56) | **2.47E-04** |
| CpG_2 | 0.46(0.32-0.58) | 0.38(0.28-0.47) | 1.33(1.11-1.58) | **0.002** |
| CpG_3 | 0.15(0.06-0.26) | 0.12(0.04-0.18) | 1.36(1.05-1.75) | **0.019** |
| CpG_4 | 0.25(0.13-0.39) | 0.21(0.08-0.30) | 1.31(1.08-1.59) | **0.007** |
| **375 BTN vs. 280 PTC (Stage I)** |  |  |  |  |
| CpG_1 | 0.48(0.36-0.65) | 0.33(0.21-0.47) | 1.51(1.37-1.65) | **3.32E-18** |
| CpG_2 | 0.46(0.32-0.58) | 0.33(0.23-0.44) | 1.53(1.38-1.70) | **2.77E-15** |
| CpG_3 | 0.15(0.06-0.26) | 0.08(0.03-0.17) | 1.59(1.38-1.85) | **7.14E-10** |
| CpG_4 | 0.25(0.13-0.39) | 0.14(0.07-0.25) | 1.48(1.32-1.66) | **7.71E-12** |
| **375 BTN vs. 44 PTC (Stage II)** |  |  |  |  |
| CpG_1 | 0.48(0.36-0.65) | 0.43(0.25-0.51) | 1.32(1.11-1.56) | **0.001** |
| CpG_2 | 0.46(0.32-0.58) | 0.37(0.25-0.50) | 1.33(1.10-1.61) | **0.003** |
| CpG_3 | 0.15(0.06-0.26) | 0.10(0.02-0.15) | 1.63(1.18-2.27) | **0.003** |
| CpG_4 | 0.25(0.13-0.39) | 0.17(0.07-0.25) | 1.38(1.10-1.73) | **0.006** |

<sup>s</sup>Logistic regression with adjustment for covariates. Significant *P* values are in bold.
Abbreviation: BTN, benign thyroid nodule; PTC, papillary thyroid cancer; IQR, interquartile range.

**Table 5.** Correlation between *RUNX1* methylation and the clinical characteristics of PTC

| Clinical characteristics | Group (n) | Median of methylation levels |  |  |  |
| --- | --- | --- | --- | --- | --- |
|  |  | CpG_1 | CpG_2 | CpG_3 | CpG_4 |
| Tumor length (cm) | < 1.50 (175) | 0.33(0.21-0.46) | 0.33(0.23-0.44) | 0.07(0.03-0.17) | 0.13(0.06-0.24) |
|  | ≥ 1.50 (149) | 0.35(0.22-0.48) | 0.34(0.24-0.46) | 0.11(0.04-0.17) | 0.16(0.08-0.28) |
|  | *P*<sup>*</sup> value | 0.655 | 0.523 | 0.144 | 0.161 |
| Tumor size | T1 (271) | 0.33(0.21-0.46) | 0.32(0.23-0.44) | 0.08(0.03-0.16) | 0.14(0.07-0.24) |
|  | T2&3&4 (53) | 0.37(0.25-0.51) | 0.38(0.28-0.47) | 0.12(0.04-0.18) | 0.21(0.08-0.30) |
|  | *P*<sup>*</sup> value | 0.177 | 0.073 | 0.209 | 0.085 |
| Lymph node involvement | N0 (138) | 0.32(0.20-0.46) | 0.33(0.23-0.45) | 0.08(0.03-0.16) | 0.14(0.07-0.24) |
|  | N1 (185) | 0.35(0.23-0.49) | 0.34(0.24-0.44) | 0.09(0.03-0.17) | 0.15(0.08-0.27) |
|  | *P*<sup>*</sup> value | 0.117 | 0.073 | 0.209 | 0.085 |
| Tumor stage | Stage I (280) | 0.33(0.21-0.47) | 0.33(0.23-0.44) | 0.08(0.03-0.17) | 0.14(0.07-0.25) |
|  | Stage II (44) | 0.43(0.25-0.51) | 0.37(0.25-0.50) | 0.10(0.02-0.15) | 0.17(0.07-0.25) |
|  | *P*<sup>*</sup> value | 0.068 | 0.119 | 0.782 | 0.761 |

All *P*<sup>\*</sup> values were calculated by Mann-Whitney U test. Significant *P* values are in bold.
Abbreviation: PTC, papillary thyroid cancer.

### Validation of the clinical value of *RUNX1* hypomethylation in distinguishing early-stage PTC from BTN

To evaluate the potential clinical application of *RUNX1* hypomethylation as a biomarker of PTC cases, receiver operating characteristic (ROC) curve analysis was performed and binary logistic regression analysis was conducted. First, the clinical efficacy of *RUNX1* hypomethylation was assessed in two independent case-control studies (Validation Ⅰ and Validation Ⅱ), and the area under the ROC curve (AUC) was 0.74 (95% CI: 0.69-0.80) and 0.79 (95% CI: 0.75-0.84), respectively (Figure 4A and 4B). Next, Validation Ⅰ and Validation Ⅱ were combined and stratified to investigate the effect of age, gender and clinical characteristics on the clinical value of *RUNX1* hypomethylation. The results showed that the AUC (95% CI) was 0.78 (0.71-0.85) in subjects < 55 years old, and 0.76 (0.72-0.80) in subjects ≥ 55 years old (Figure 4C and 4D). The AUC (95% CI) was 0.72 (0.64-0.80) and 0.77 (0.73-0.81) in male and female subjects, respectively (Figure 4E and 4F). Furthermore, the AUC (95% CI) was 0.80 (0.75-0.85) in BTN subjects and PTC cases with T1, and 0.79 (0.75-0.82) in BTN subjects and PTC cases at Stage Ⅰ (Figure 4G and 4H). Together, the AUC results demonstrated that the *RUNX1* hypomethylation based on FFPE tissue samples has high credibility and accuracy in distinguishing early-stage PTC from BTN, especially for younger women and PTC cases with T1 or at Stage Ⅰ.

**Figure 4.**
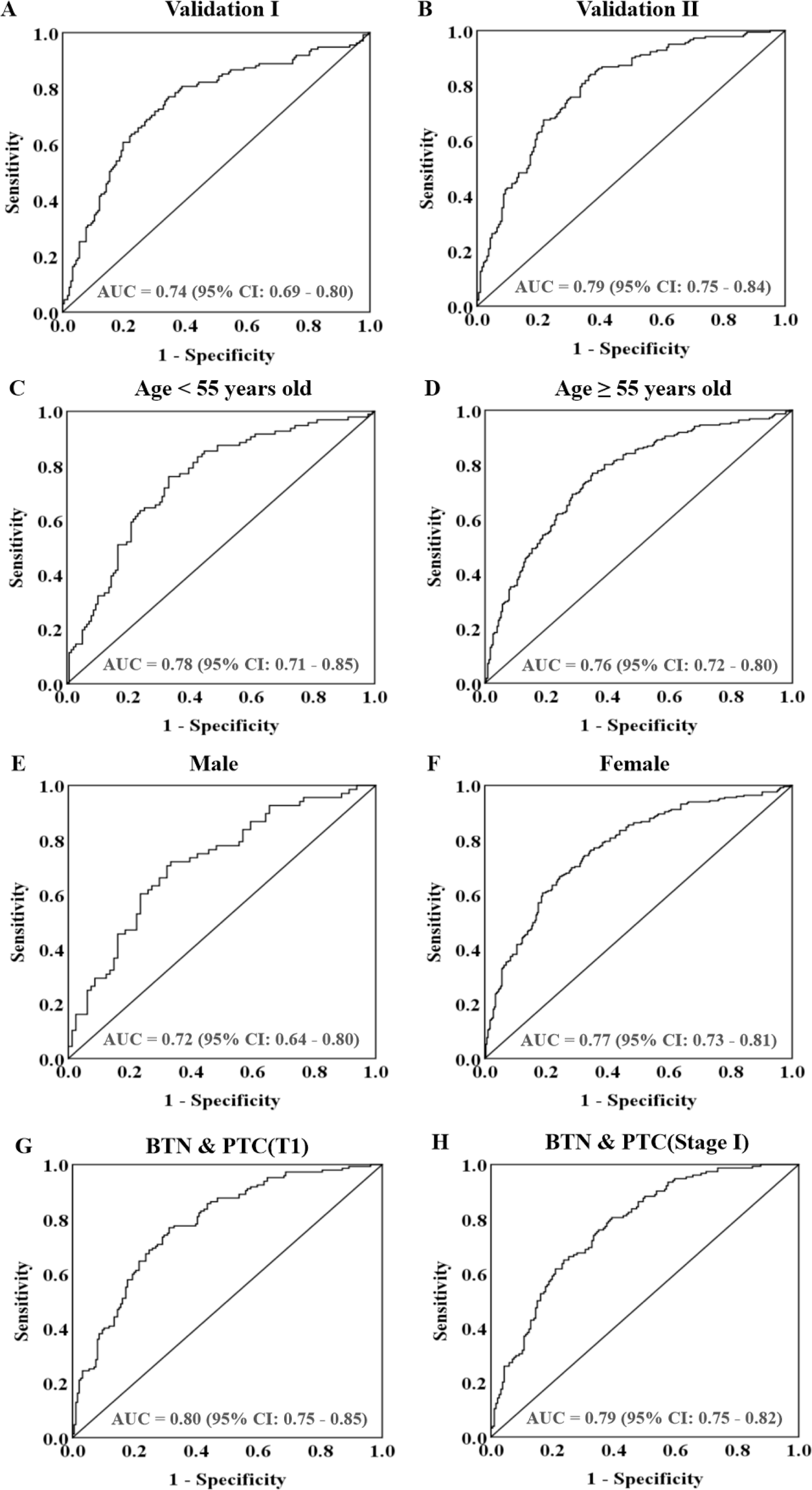
The clinical value of *RUNX1* methylation in distinguishing early-stage PTC cases from BTN patients. The methylation levels of four CpG sites within *RUNX1* gene were generated a prediction probability, (A-B) ROC curve analyses for the discriminatory power of *RUNX1* prediction probability to distinguish PTC cases from BTN subjects in Validation Ⅰ (A) and Validation Ⅱ (B). In addition, Validation Ⅰ and Validation Ⅱ were combined and stratified by the age of 55 years old, gender, tumor size or tumor stage, respectively. (C-H) ROC curve analyses for the discriminatory power of *RUNX1* prediction probability to distinguish PTC cases from BTN subjects in age < 55 years old group (C), age ≥ 55 years old group (D), male group (E), female group (F), tumor size group (G) and tumor stage group (H), respectively. All the above 95% confidence interval (CI) of AUC were calculated by logistic regression with covariates-adjusted.

## Discussion

In clinical practice, it is necessary to explore objective and stable biomarkers for accurate diagnosis of thyroid nodules, which can not only avoid over-diagnosis and over-treatment in BTN patients, but also provide timely treatment for patients with nodules of malignant potential. Recently, the relationship between DNA methylation and thyroid cancer has been studied intensively and attracted wide attention[17]. However, most reports have focused on cancerous and para-cancerous tissues, while only a few studies involve the identification of benign and malignant thyroid tumors, mostly based on candidate genes[24, 25]. In addition, these studies have shortcomings such as small sample size and poor systematic research content. The present study systematically investigated the value of tissue DNA methylation in the differential diagnosis of thyroid nodules in Chinese population, thereby providing new evidence for potential clinical application of methylated genes as cancer biomarkers. Furthermore, this study has reliable methods, rigorous design and sufficient sample size. The combined analysis of 850K beadchip array and RNA-Sequencing in the fresh frozen tissue samples of early-stage PTC cases and BTN subjects revealed a strong negative correlation between reduced DNA methylation and increased mRNA of *RUNX1*. Subsequently, *RUNX1* hypomethylation in PTC was validated in two independent studies with FFPE samples from a total of 699 BTN and PTC subjects using mass spectrometry.

RUNX proteins are a family of transcription factors with highly conserved sequences, which are involved in cell growth, proliferation and differentiation[26]. *RUNX1* (runt related transcription factor 1) is also known as acute myeloid leukemia 1 (*AML1*), and its abnormal expression or mutation can lead to human leukemia[27]. Expression of RUNX1 is regulated by DNA methylation, post-translational modifications (e.g., phosphorylation, acetylation, and ubiquitination)[28]. First identified in acute myeloid leukemia, *RUNX1* has also been found to play contrary roles in different solid tumors. For example, *RUNX1* acts as a tumor suppressor gene in hepatocellular carcinoma and gastric cancer, but as an oncogene in non-small cell lung cancer and endometrial cancer[29–32]. *RUNX1* is mainly involved in the regulation of TGF-β, WNT and BMP (bone morphogenetic protein) signaling pathways[33]. Previous studies have shown that abnormal *RUNX1* methylation was implicated in tumor formation and malignant progression. For example, hematopoietic stem cells acquire survival advantage by *RUNX1* hypomethylation in familial leukemia[34]; abnormal methylation in *RUNX* family is negatively correlated with immune cell infiltration in breast cancer[35]. So far, abnormal methylation of *RUNX1* has not been reported in thyroid related diseases. Here, we found *RUNX1* methylation was significantly reduced and its mRNA was significantly increased in early-stage PTC cases, suggesting that *RUNX1* may act as an oncogene in the progression of PTC.

DNA methylation is an early and concomitant event during cancer progression[36]. It has been well documented that cancer is a disease closely associated with aging and may have similar abnormalities in DNA methylation, mainly including hypomethylation in oncogene promoter regions or widespread hypermethylation in gene body[16, 37]. Due to the influence of sex hormones, there are inherent differences in DNA methylation at CpG sites in certain genes between males and females[38, 39]. In China, the incidence of TC increases significantly along with aging, and reaches its peak at 55 years old, and the incidence is three times higher in women than in men[40, 41]. Therefore, both age and gender are key factors in the formation and malignant progression of TC[42]. The present study revealed that the association between *RUNX1* hypomethylation and PTC cases was more obvious in subjects younger than 55 years and females in age- and gender-stratified analyses, suggesting that *RUNX1* hypomethylation was a very important risk factor for PTC cases in young woman.

Clinically, tumor length, tumor size, tumor stage and lymph nodes involvement are strongly associated with malignant progression and poor prognosis of PTC patients[43, 44]. In this study, no significant correlation was observed between *RUNX1* methylation and malignant progression of PTC. Interestingly, we found differences in *RUNX1* methylation between PTC cases with T1 or at Stage Ⅰ and BTN subjects were more significant than those between PTC cases with T2&3&4 or at Stage Ⅱ and BTN subjects. Besides, *RUNX1* methylation achieved good accuracy in differentiating early-stage PTC cases with T1 or at Stage Ⅰ from BTN subjects. Although these results might be less credible due to the small sample size of PTC patients with larger tumor size or at advanced stages, our findings suggested that the modulation of *RUNX1* methylation may play different roles in the initiation and progression of PTC, and further indicated the complex epigenetic regulation in cancer. The association between *RUNX1* hypomethylation and PTC malignant progression needs further evaluation in future studies involving more subjects.

Both benign and malignant thyroid tumors have abnormal cell proliferation, DNA mutations or epigenetic changes, and large tumors are often accompanied by abundant angiogenesis and energy consumption[45]. Nevertheless, benign tumors don’t invade neighboring tissue or metastasize and can be safely monitored with no or minimal treatment, whereas malignant tumors are usually aggressive and have a chance of recurrence and metastasis after resection of the focus, seriously threatening the life and health of patients[46, 47]. Therefore, accurately identifying malignancy of thyroid tumors, especially those with indeterminate cytology, is essential for avoiding overtreatment and controlling tumor metastasis, which could benefit from supportive biomarkers. In this study, we identified *RUNX1* methylation with high credibility and good accuracy in differentiating early-stage PTC cases from BTN subjects, suggesting it as a potential biomarker to aid the diagnostic process[47, 48]. Future studies with larger sample size are warranted to further evaluate the clinical value of *RUNX1* methylation in assessing different subtypes and the aggressiveness of thyroid cancer. The function and underlying molecular mechanisms of *RUNX1* hypomethylation and overexpression in the malignant progression of thyroid cancer need further investigation[49]. Our results demonstrate a new tumor-driven indicator, and provide a novel insight into the etiology of PTC.

## Conclusion

Taken together, this study revealed significant hypomethylation and overexpression of *RUNX1* in early-stage PTC cases compared to BTN subjects in genome-wide study, and further validated the alter *RUNX1* methylation in two independent case-control studies by mass spectrometry. Our findings suggested that *RUNX1* methylation might be used as a novel biomarker to distinguish malignant and benign thyroid nodules.

## Statements and Declarations

### Funding

This study was financed by the research funding of Nanjing Medical University and the research funding of the Jiangsu Province.

### Competing interests

The authors have no relevant financial or non-financial interests to disclose.

### Data availability statement

The authors affirm that all data necessary for confirming the conclusions of this article are represented fully within the article and its tables and figures.

### Authors’ contributions

Junjie Li, Chenxia Jiang and Rongxi Yang conceived and designed this study; Junjie Li and Yifei Yin wrote the main manuscript and prepared figures and tables; Junjie Li, Haixia Huang and Mengxia Li conducted major experiments; Hong Li and Minmin Zhang were responsible for data interpretation and analysis; Junjie Li drafted and Rongxi Yang revised the manuscript. All authors reviewed the manuscript.

